# High-Performance Image-Based Measurements of Biological Forces and Interactions in a Dual Optical Trap

**DOI:** 10.1101/430686

**Authors:** Jessica L. Killian, James T. Inman, Michelle D. Wang

## Abstract

Optical traps enable nanoscale manipulation of individual biomolecules while measuring molecular forces and lengths. This ability relies on the sensitive detection of optically trapped particles, typically accomplished using laser-based interferometric methods. Recently, precise and fast image-based particle tracking techniques have garnered increased interest as a potential alternative to laser-based detection, however successful integration of image-based methods into optical trapping instruments for biophysical applications and force measurements has remained elusive. Here we develop a camera-based detection platform that enables exceptionally accurate and precise measurements of biological forces and interactions in a dual optical trap. In demonstration, we stretch and unzip DNA molecules while measuring the relative distances of trapped particles from their trapping centers with sub-nanometer accuracy and precision, a performance level previously only achieved using photodiodes. We then use the DNA unzipping technique to localize bound proteins with extraordinary sub-base-pair precision, revealing how thermal DNA fluctuations allow an unzipping fork to sense and respond to a bound protein prior to a direct encounter. This work significantly advances the capabilities of image tracking in optical traps, providing a state-of-the-art detection method that is accessible, highly flexible, and broadly compatible with diverse experimental substrates and other nanometric techniques.

## Introduction

Optical traps (OTs) have made indelible contributions to our understanding of nanoscale biophysical processes through their powerful abilities to measure forces and distances while actively manipulating individual biomolecules^1^. Their determination of molecular forces with sub-piconewton accuracy on sub-millisecond timescales relies on a fundamental measurement: the displacement of a trapped microsphere (bead), a reporter particle attached to a biomolecule of interest, from the trap’s center. Traditionally, this quantity is obtained through the technique of back-focal-plane interferometry (BFPI)^2^, which uses a photodiode to measure the deflection of the trapping laser that occurs when a bead is displaced from the trap’s center. The exceptional bandwidth, precision, and accuracy provided by BFPI has enabling such feats as the observations of single base-pair steps by translocating motor proteins^3,4^, and the detailed characterization of histone-DNA interactions with near base pair resolution^5^.

Camera-based particle tracking is an appealing alternative to BFPI due to its ability to make direct measurements of positions and distances in the sample plane, its flexibility, and its relative ease of implementation on a commercial microscope. Over the past two decades, the use and efficacy of image tracking in scientific instruments has flourished in direct response to a remarkable revolution in digital camera technology that has yielded dramatic improvements in data bandwidth, sensor noise, and real-time tracking techniques^6–9^. Indeed, image tracking is already widely used in the field of magnetic tweezers where the absolute positions of reporter particles are tracked with sub-nanometer precision at multi-kilohertz speeds^7,9,10^, and has also been used recently to localize optically-trapped particles in solution with similar precision and speed^11–13^. However, despite these achievements, image tracking has not yet demonstrated the ability to make accordingly accurate real-time measurements of biological forces in an OT.

This discrepancy has a subtle but important source: optical traps present unique challenges for image-based force measurements in that it is the bead’s displacement from trap center, rather than its absolute position, that is of interest. While BFPI obtains this quantity inherently and accurately in a single measurement, cameras show only the bead’s absolute position and offer no coincident measure of trap position. In practice, even the most precise pre-determined estimates of trap position in the sample plane are rendered inaccurate by laser pointing fluctuations and drift^14^, which alter trap position by tens to hundreds of nanometers over experimental timescales. Consequently, an image-based OT able to perform advanced single-molecule manipulation experiments and obtain biological measurements of the same exceptional quality as its precision photodiode-based counterparts has not yet been realized.

Here we present an adept dual optical trap and a set of camera-based detection techniques that overcome these challenges. We determine biological forces by measuring the relative displacement of the beads within our optical traps with both sub-nanometer precision *and* sub-nanometer accuracy, across a trap movement range of nearly 10 ⍰m, and at up to 10 kHz in real time using a unique and low-latency field-programmable gate array (FPGA) tracking implementation. We demonstrate our instrument’s capabilities by stretching and unzipping DNA molecules, and then use the versatile DNA unzipping technique to localize bound proteins with remarkable sub-base-pair precision. Revealed by the instrument’s exceptional data quality, we additionally find that thermal DNA fluctuations enable an approaching unzipping fork to sense and respond to a bound protein prior to a direct encounter, and model the sequence-dependence of this behavior theoretically.

## Results

### Instrument Design

Maximizing the performance of camera-based optical trap detection requires both a stable optical trap and a capable image tracking system. Our instrument (Fig. 1a, Supplementary Fig. 1, further details in the online methods and Supplementary Note 1) utilizes a dual optical trap, created by time-sharing a single laser trap between two locations in the sample plane. Because they share identical optical paths, traps generated by timesharing are remarkably self-consistent in their relative positions^15^. We use an acousto-optic deflector (AOD) to timeshare our traps at 50 kHz, providing low-noise and high-stiffness traps^16^ and allowing independent control of the trap powers and positions along a single axis.

**Figure 1.**
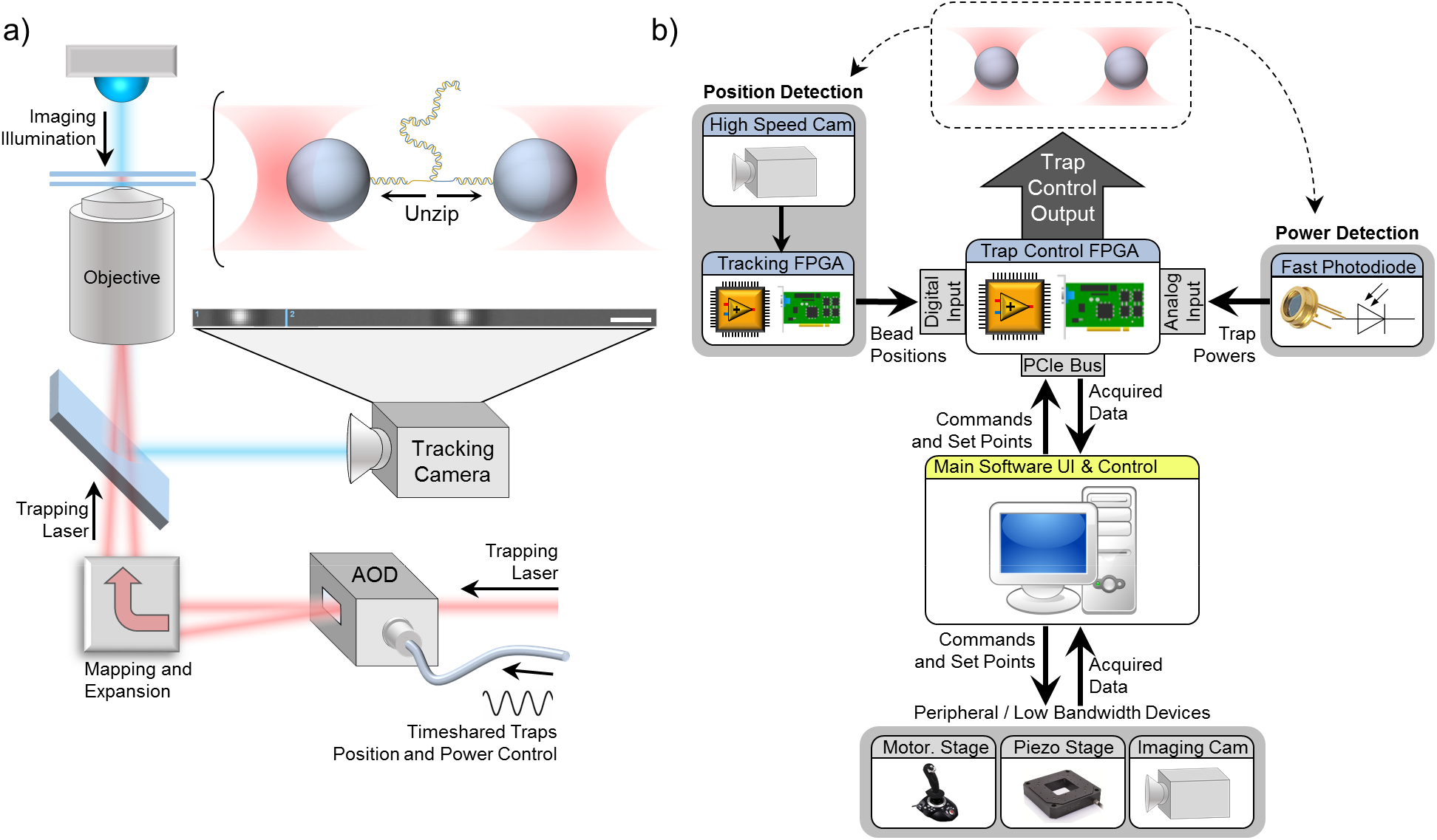
Instrument Overview. **(a)** Dual optical traps are generated by timesharing a single laser via an AOD, which controls the positions and powers of both traps in the sample plane. Bead positions are tracked in real time at up to 10 kHz using a high speed camera. The drawn scale bar is 400 nm. **(b)** The control flow of the instrument. A “trap control” FPGA generates the driving signal to control the AOD (and thus the traps) in Fig. 1a, and acquires and processes instrument data including trap powers from a high speed photodiode and bead positions from a second “image tracking” FPGA. Acquired data sampled at up to 50 kHz are streamed to the the main host software for display and writing to disk.

The AOD is driven by an RF synthesizer, which we control using an FPGA. This “trap control FPGA” acts as a centralized instrument hub, acquiring and processing instrument data, and then determining and generating the driving signal for the RF synthesizer to control the optical traps (Fig. 1b). Trap powers are measured for each trap individually by a high-speed photodiode and stabilized via feedback (Supplementary Fig. 2). Bead position detection (detailed in the following section) is performed by a high-speed camera connected directly to a second “image tracking FPGA” dedicated to tracking bead positions. Thus the system provides an embedded platform for data acquisition, feedback, and instrument control, executed with hardware-timed determinism on the FPGA’s 40 MHz clock. Custom LabVIEW 2015 software directs the top-level instrument operation.

Achieving both high-speed and precise image tracking requires a low-noise camera sensor and a bright light source to provide adequate illumination at sub-millisecond camera exposures^17^. We acquire a cropped region of interest (Fig. 1a) at up to 10 kHz using a CMOS camera illuminated by a high-powered LED. Additionally, we incorporate a 4× optical magnification on top of our 60× water-immersion objective, resulting in images scaled at 57.3 nm per pixel. A large image magnification improves tracking accuracy by reducing the relative contributions, if present, of tracking algorithm bias and camera fixed pattern noise^7^. Once acquired, image data from the camera are transferred over a Camera Link interface directly to the image tracking FPGA for processing.

### Cross-Correlation Image Tracking on an FPGA

Real-time image tracking at high camera framerates generates large amounts of data, which strains the throughput capabilities of traditional software tracking and requires a specialized solution for accelerated processing. This is typically realized by performing all or part of an image tracking algorithm on a graphics processing unit (GPU).^7,9^–12 Less commonly used, an FPGA provides a re-programmable hardware implementation of a tracking algorithm, yielding superior determinism and the low latencies required for demanding real-time control applications^18^. On the other hand, their limited physical resources and lack of native floating point support constrains their aptitude for complex tracking algorithms. Here we have achieved a fast, ultra-low-latency FPGA implementation of the precise and accurate mirror cross-correlation (MCC) image tracking algorithm^19^ (Fig. 2a).

**Figure 2:**
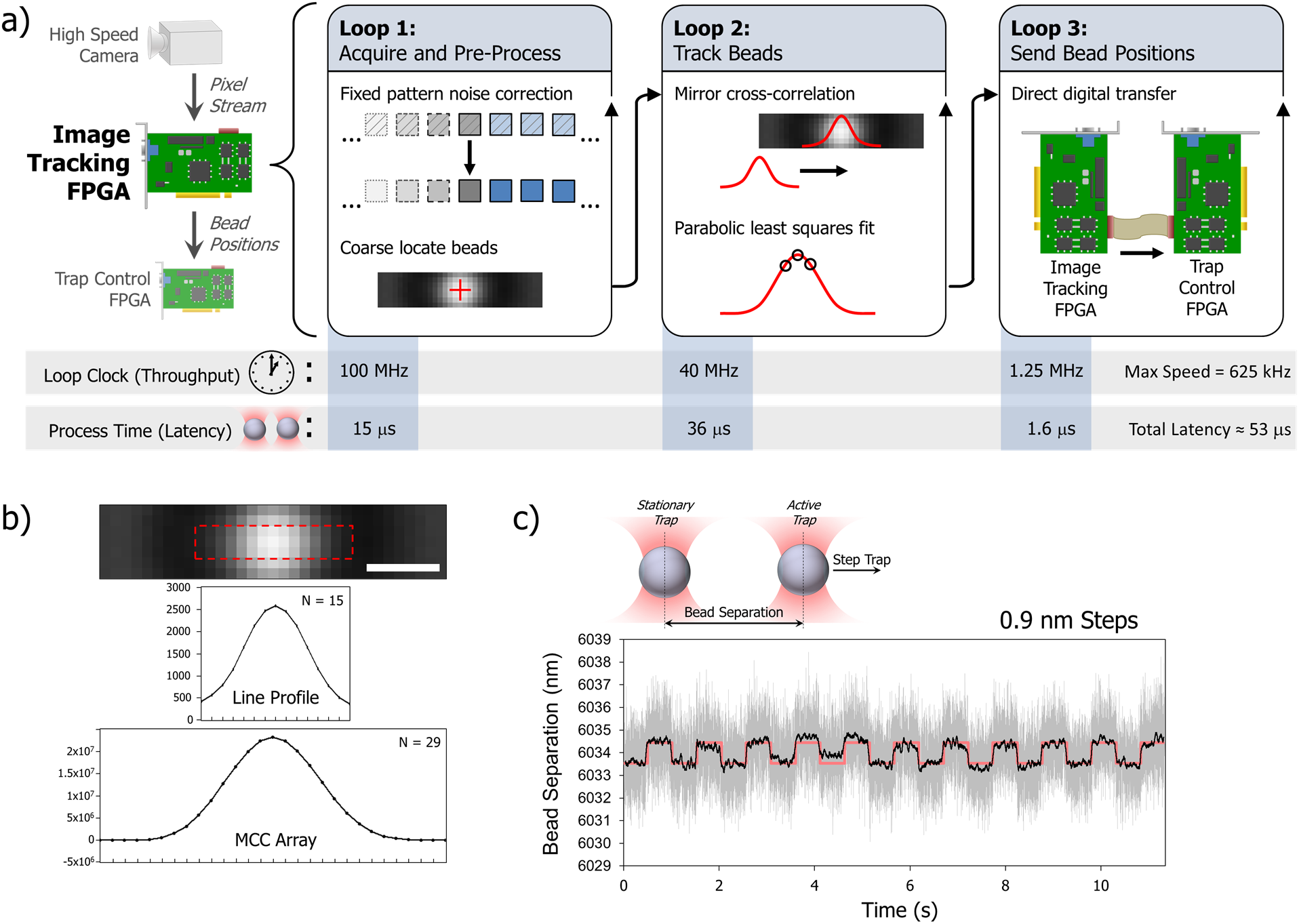
FPGA image tracking and results. (**a**) FPGA image tracking using a cross-correlation algorithm is implemented in three independent loops, the first for image pre-processing, the second for bead tracking, and the third for data transfer. (**b**) A cropped 10-bit image of an 800 nm polystyrene bead taken from experimental data, and its corresponding line profile is shown below. The results of cross-correlating the line profile with its mirror image (“MCC Array”) are shown in the bottom panel. (**c**) With a bead in each trap, we held one trap stationary and stepped the other trap in increments of 0.9 nm while measuring the bead separation. The raw data (2.5 kHz) is shown in gray, while the black trace is filtered to 250 Hz. The trap control driving signal is shown in light red.

Image pixels are streamed directly from the camera to the image tracking FPGA, where we first correct for the camera’s fixed pattern noise and then determine the positions of the bead centers. We track beads in one dimension (along the tethering axis), which is sufficient to reach the thermal limit of detection in dual optical traps^20^. The bead positions are then sent to the trap control FPGA over directly-wired digital lines. Overall, the system permits tracking and transfer of the dual bead positions at up to 625 kHz (though our camera configuration limits us to 10 kHz acquisition rates), with a latency between the finished camera exposure and the completed transfer of both tracked bead positions of about 50 μs. The tracking exhibits both high precision and high accuracy; the variance in the tracked position of a bead fixed firmly to a coverslip stays below 0.2 nm^2^ through 10 kHz acquisition rates, and a histogram of sub-pixel bead positions shows minimal bias (Supplementary Fig. 3). Sub-nanometer changes in the separation between two strongly trapped 800‐nm beads (Fig. 2c) are clearly visible.

### Accurate Force Measurements in Mobile Traps

For small bead displacements from the trap centers, force in a dual optical trap can be expressed as *F* = *K*_eff_ Δ*X*_bead_ where *K*_eff_ is the effective trap stiffness of the two traps and Δ*X*_bead_ is the average displacement of the two trapped beads relative to their respective trap centers (Fig. 3a, Supplementary Note 2). While image tracking is suited to the task of determining absolute bead positions in the sample plane, the large uncertainty in the concurrent trap positions ultimately limits the accuracy of individual trap force measurements. However, while the position of an optical trap in the sample plane fluctuates and drifts substantially, the separation between two time-shared traps may be much more stable^20^ (Supplementary Fig. 4). Our technique for measuring force (Fig. 3a) leverages the greatly enhanced stability of timeshared dual optical traps by forgoing the use of absolute positions altogether and relying on an expression of Δ*X*_bead_ that uses only separations.

**Figure 3:**
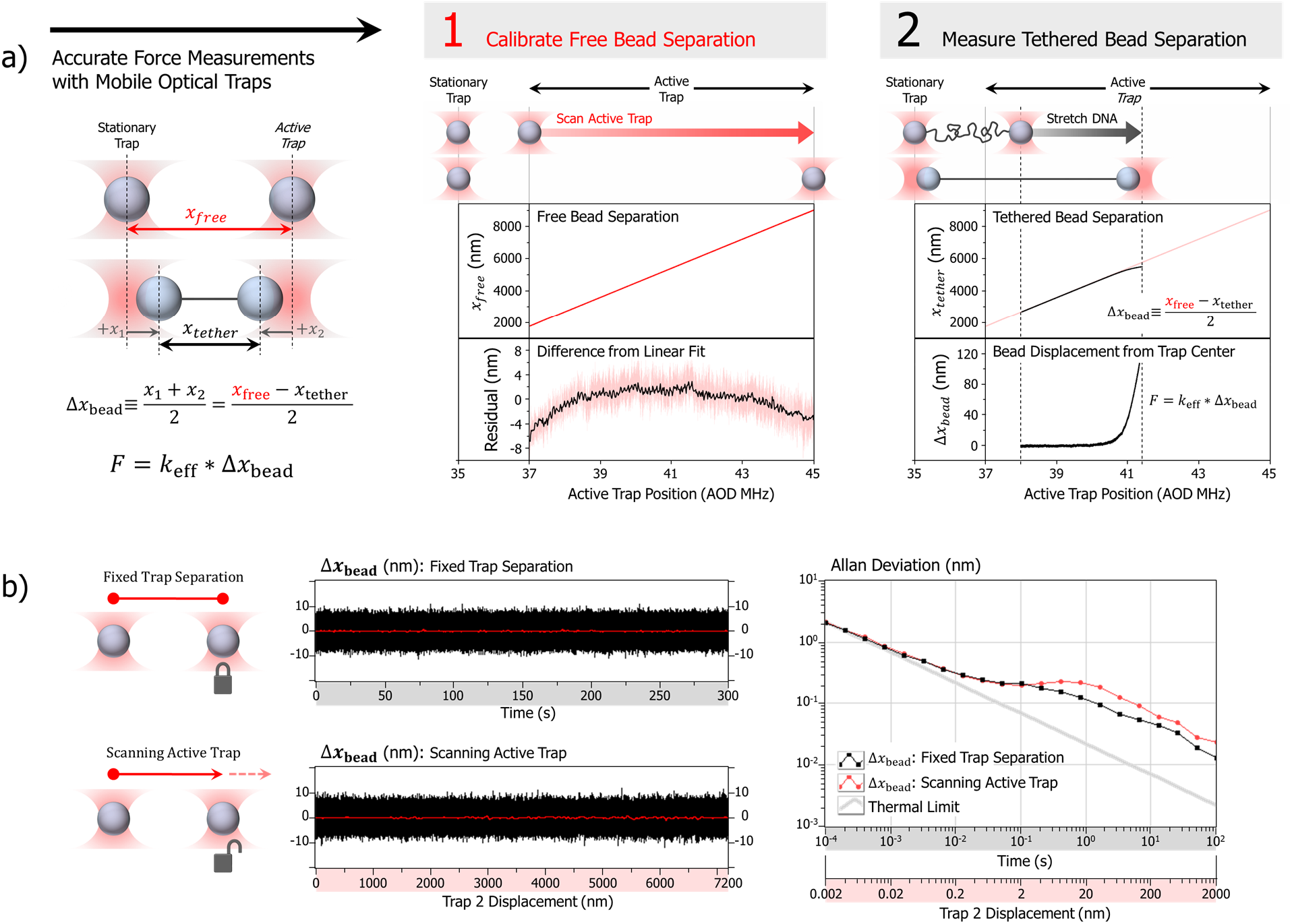
Accurate image-based force measurements with mobile traps. (**a**) Force in a dual optical trap can be expresses in terms of the relative trap separation coordinates *X*_free_ and *X*_tether_. *X*_free_ versus trap position is first measured and stored as a LUT on the trap control FPGA prior to an experiment. Half the difference between *X*_free_ indexed from this calibration LUT, and *X*_tether_ measured during an experiment, yields Δ*X*_bead_ for each image frame. **(b)** The Allan Deviation (ADEV) of the Δ*X*_bead_ signal for stationary traps was compared to ADEV of the Δ*X*_bead_ signal as the active trap was scanned across its mobile range, revealing the accuracy of Δ*X*_bead_ measurements in mobile traps.

We begin by calibrating the separation of the traps. Holding an untethered (free) bead in each of the two traps, we track the separation between them (*X*_free_) as we maintain one trap stationary and step the second “active” trap across its remaining mobile range. The free bead separation, reflecting the absence of an externally applied force, represents the center-to-center distance between the two traps. The measured trap separation at each AOD frequency of the active trap (which drives its position in the sample plane), is sent to the trap control FPGA as a look-up-table (LUT) prior to an experiment. As Figure 3a illustrates, the use of a LUT in this process is critical to measurement accuracy due to the nonlinearity of the trap response to its control signal.

When experiments with a biological substrate are performed, we again fix the stationary trap, then manipulate the sample via movement of the active trap while measuring the separation between the two beads (*X*_tether_). The AOD frequency of the active trap serves to index the LUT of trap separation (*X*_free_). Half of the difference between *X*_free_ and *X*_tether_ represents the average displacement of the beads in their respective traps for that image frame, Δ*X*_bead_. The accuracy of this technique relies on the stability of the trap separation calibration afforded by the time-shared dual optical traps, which confer the repeatability required for a robust measurement of force. We characterize this by obtaining the Allan Deviation (ADEV)^14^ of Δ*X*_bead_ for two free beads as the active trap is scanned across its full movement range, and comparing it to that of stationary traps over a comparable time period (Fig. 3b). Inaccuracies in the LUT calibration of *X*_free_ will manifest as non-physical fluctuations in the measured value of Δ*X*_bead_ as the active trap is scanned, and a consequent increase in the ADEV of the moving traps relative to the stationary traps. The results show that this technique enables us to obtain Δ*X*_bead_ with sub-nm accuracy across the full mobile trap separation range.

### Stretching and Unzipping DNA

The ability to make sensitive measurements of biological forces and distances is the cornerstone of single-molecule optical trapping studies. Thus to demonstrate our instrument’s ability to acquire high-quality data in single-molecule manipulation experiments, we begin with two classic and well-characterized optical trapping assays: stretching and unzipping DNA. Throughout both experiments, *K*_eff_ was held at ^~^0.3 pN/nm, thus small inaccuracies in the measure of bead displacement would have a large effect on the measured force.

The force-extension relationships from stretching 12 individual duplex DNA molecules over the course of an hour are shown in Figure 4a, along with the theoretical prediction (not a fit)^21^. These data were aligned in extension via small (<10 nm) global offsets, accounted for by variations in bead diameters (Supplementary Fig. 5), but are unaligned in force. Next, we used our instrument to unzip DNA, which describes the mechanical disruption of the duplex DNA into its two constituent strands^22^. In Figure 4b we show a zoomed in region of raw, unaligned data obtained from unzipping and re-zipping a single tether 3 times over a period of about 3 minutes. As the molecule is unzipped, the force is observed to vary according to the underlying base pair sequence, for which the theoretical prediction is shown in red^22^. The data for both stretching and unzipping exhibit exceptional agreement with respect to the theoretical predictions, highlighting the accuracy of the measurement technique. Also remarkable is the trace-to-trace consistency, highlighting the stability and low drift of the instrument over long experiment timescales.

**Figure 4:**
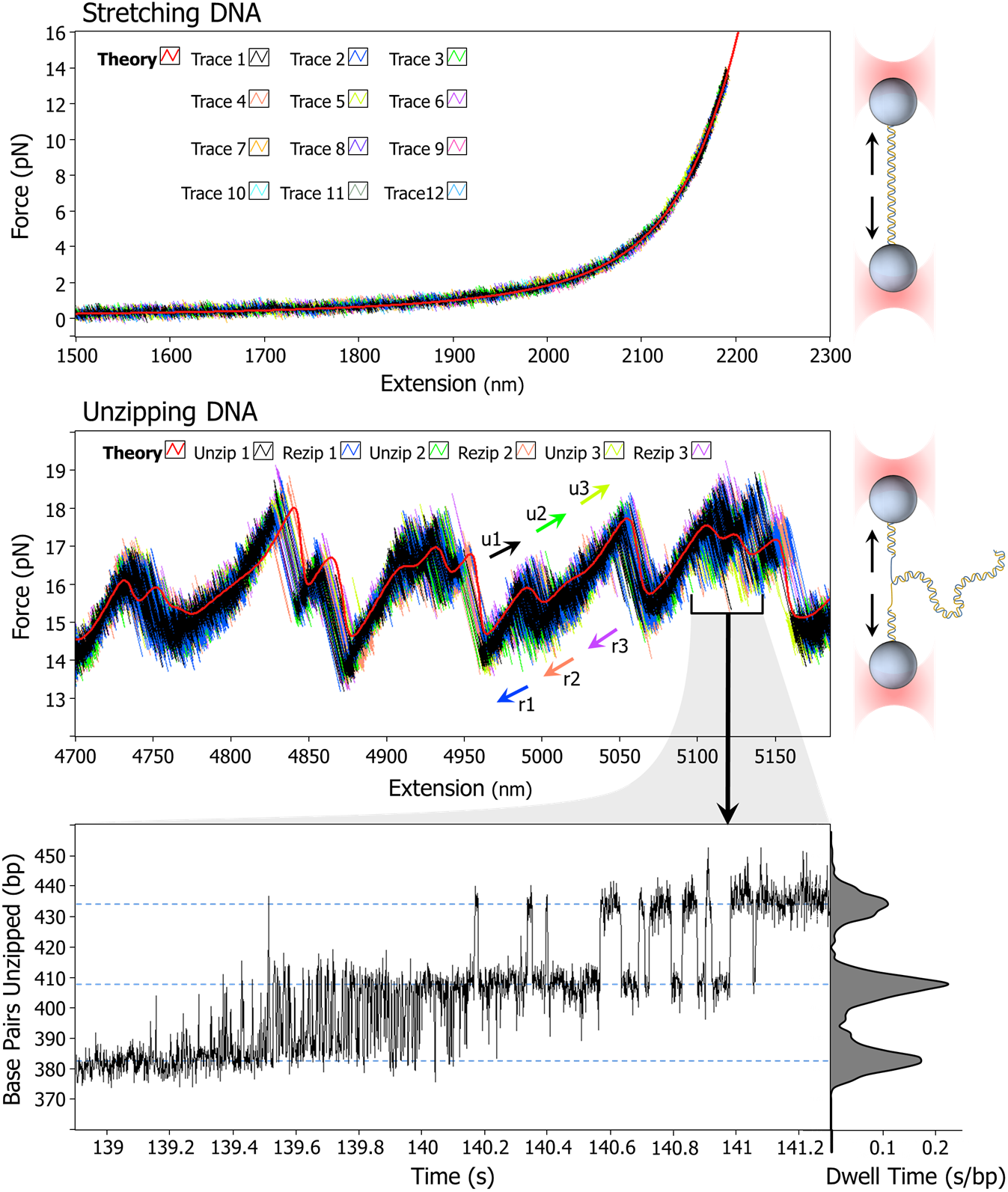
Stretching and unzipping naked DNA. (**a**) The measured force-extension relationships for twelve different DNA molecules are shown, along with the theoretical prediction^21^ (not a fit). **(b)** A single DNA molecule was unzipped until just before strand dissociation and then rezipped. This process was performed three consecutive times over a period of about 3 minutes. **(c)** A portion of the data in Figure 4b was converted to the number of base pairs unzipped versus time. Rapid transitions on millisecond timescales between discrete unzipped states are visible. A dwell time histogram is also shown.

In Figure 4c we examine a portion of the unzipping data from Figure 4b which has been converted to display the number of base pairs unzipped as a function of time. With this presentation, we show sufficient bandwidth and precision to resolve discrete hopping fluctuations between nearby unzipping states on millisecond timescales. This behavior is not unexpected; thermally-driven “breathing” -- rapid and spontaneous opening and closing of DNA base-pairing interactions -- is highly characteristic of DNA fork junctions and is thought to play important regulatory roles in *in vivo^23,24^*.

### Fork Breathing Fluctuations Modulate Protein Unzipping

Unzipping DNA reveals a detailed and distinct force signature shaped by the underlying energy landscape of its sequence, and thus unzipping also serves as a powerful tool for probing and localizing protein-DNA interactions^5,25–27^. A bound protein presents a roadblock which increases resistance to unzipping, resulting in a detectable rise in force above the naked DNA baseline when the unzipping fork encounters the protein. The location of the force rise thereby communicates the location of the bound protein on its DNA substrate. To investigate the performance of our instrument in an advanced setting, we unzipped through proteins bound to sequence-specific sites on DNA molecules in order to localize their interactions. Interestingly, enabled by the instrument’s exceptional data quality, our experiments uncovered unexpected insights into the effects of DNA fork breathing fluctuations on the protein unzipping process.

Figure 5 shows two sets of data obtained by unzipping through the bound restriction enzyme HincII. Though the protein and its recognition sequence are the same for both data sets, the DNA sequences preceding the binding sites are different. Against the backdrop of the naked DNA unzipping theory, the bound protein produces an unmistakable vertical force rise, occurring when the progression of the unzipping fork is impeded by the tightly-bound protein (Figs. 5a, 5b). Surprisingly, the data also reveal distinct deviations from the naked DNA unzipping theory beginning well before the expected vertical force rise (Figs. 5c, 5d). The correct interpretation of these data was not immediately apparent, due to several physical phenomena that could plausibly produce such effects, including protein sliding, protein remodeling, weak protein-DNA interactions outside the main binding region, and even instrument miscalibration. Ultimately, we determined this phenomenon to be consistent with just a single strong protein-DNA interaction, located at the position of the vertical force rise.

**Figure 5:**
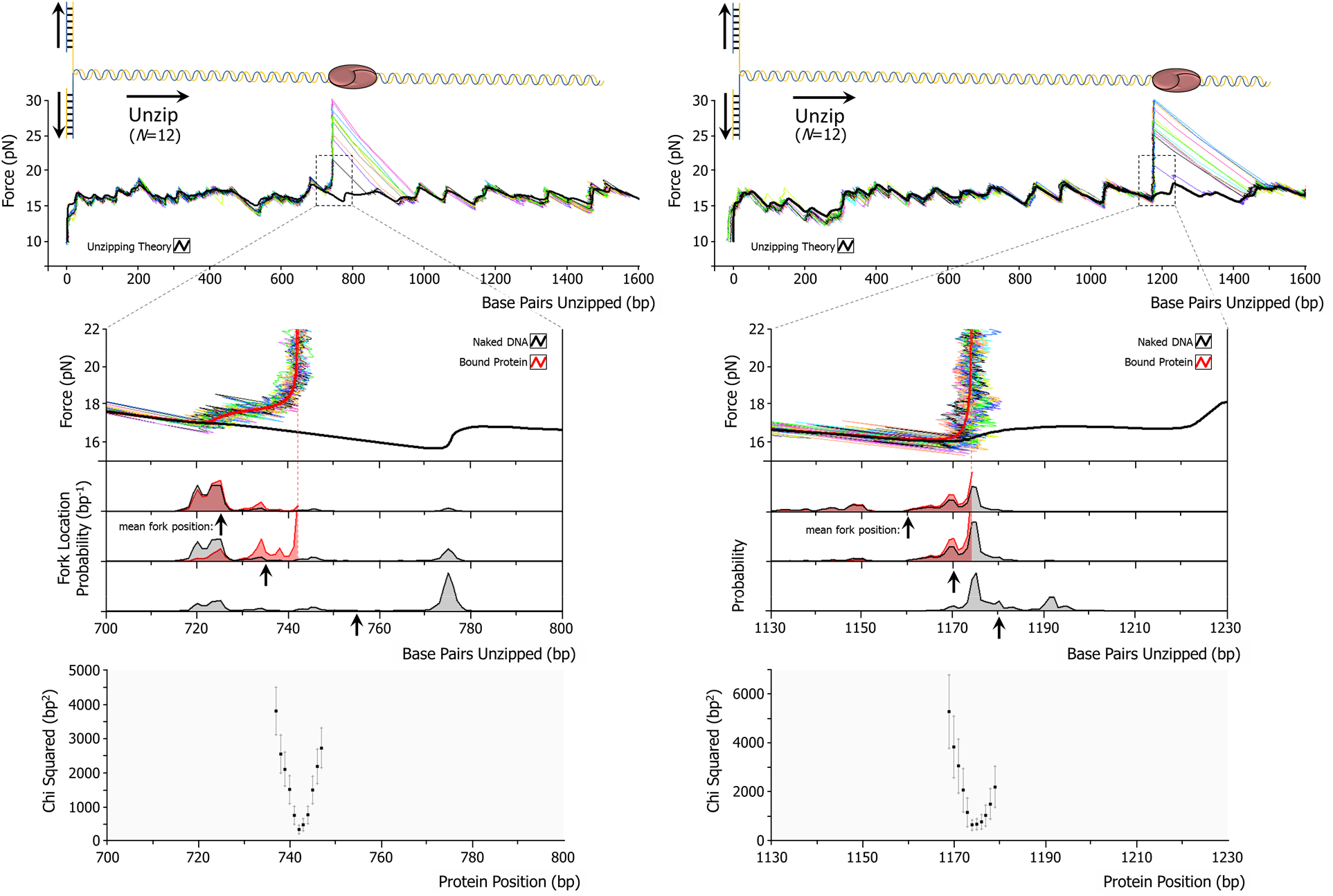
A bound protein modulates unzipping at a distance. (**a, b**) The restriction enzyme HincII was bound to a DNA template and unzipped from either direction. **(c,d)** Featured deviations from the naked unzipping theory (black) are observed as the protein is unzipped. This was predicted well by our unzipping theory (red) by placing an infinite energy barrier at a single base in the sequence. Below the data is the calculated likelihood for the fork to reside at a given sequence position with (red) and without (black) a protein bound at the dashed red line, as the mean unzipping fork position (black arrow) progresses down the DNA template. A force rise is seen once the protein perturbs the distribution of states the fork is able to explore through thermal fluctuations, which varies with the DNA sequence. (**e, f)** Theoretical protein unzipping signatures were generated for a protein bound at integer base positions along the DNA sequence, and the *χ*^2^ value of our measured data against each separate prediction was determined and fit with a parabola to extract the protein position with sub-bp precision.

How does the protein affect the unzipping force many bases upstream of its binding location? The mechanism lies in the thermally-driven breathing dynamics of the unzipping fork, which causes its location at a given value of DNA extension to fluctuate about its mean position. Consequently, both the average unzipping force and the average fork position at any given time reflect the cohort of unzipping states the fork is able to explore (Figs 5c, 5d), which is strongly influenced by the underlying DNA sequence. A force rise is induced by a bound protein when it inhibits the extent of these fork fluctuations. In Figure 5c, this occurs well in advance of the bound protein due to the large amplitude of the fluctuations through its AT-rich binding vicinity. In contrast, in Figure 5d the protein’s binding sequence is located within a GC-rich region, which itself already restricts the fluctuations of an upstream unzipping fork from progressing into the downstream DNA. Accordingly, the presence of the protein does not perturb the native unzipping behavior until the fork is immediately adjacent to the protein, resulting in a relatively sharp and abrupt transition into the protein’s vertical force rise.

Thus we find that the unzipping signature of a DNA-bound protein is highly sensitive to the DNA sequence preceding the protein binding site, via the protein’s ability to perturb the extent of the breathing fluctuations of an approaching unzipping fork. To predict the unzipping fork’s behavior using our theoretical model of unzipping, we simulate a bound protein as an infinite energy barrier at a defined sequence location such that the unzipping fork cannot progress beyond that point. The insight this provides is critical to interpreting protein unzipping data below ^~^20 pN, in order to differentiate weaker protein-DNA interactions at a given sequence location from a strong barrier further downstream. Additionally, the sensitive sequence-dependence of the low-force unzipping signature contains valuable information on the protein’s location. To take advantage of this, we generated unzipping theory curves for a protein located at successive sequence locations in the vicinity of binding site, and determined the *χ*^2^ value of our data against each theoretical protein signature (Figs. 5e, 5f, Supplementary Fig. 6). The sub-bp minimum of the *χ*^2^ versus protein position plot was determined by parabolic fit, allowing us to localize a protein-DNA interaction with high accuracy and precision, even when high force data were unavailable.

Breathing fluctuations are an intrinsic feature of single-to-double stranded DNA fork junctions both *in vitro* and *in vivo*^28^. Within the cell, these junctions occur at replication forks where, instead of mechanical unzipping, DNA is unwound by helicases. In this process, breathing fluctuations are believed to play an important regulatory roll by influencing helicase movement into the replication fork^24^. Our data suggest that, additionally, thermal fork fluctuations may allow an approaching *in vivo* unzipping fork to sense the presence of a bound protein well in advance of a direct encounter. The sequence-dependent slowing of a replication fork approaching a bound protein could potentially facilitate early information transfer before contact is made, e.g. by enhancing recruitment and binding of other proteins to assist in the roadblock’s removal. This presents a distinct mechanism for “action at a distance”^29^, by which a protein at one location on a DNA sequence communicates information to a protein some distance away.

### Unzipping Footprints of DNA-Bound Proteins

With this knowledge of protein unzipping behavior in hand, we are well-positioned to probe the interaction landscape of a bound protein. We unzipped three type II restriction enzymes, each with a unique binding site along a DNA sequence, from either direction (Fig. 6), then used the *χ*^2^ analysis technique to determine the locations of the interactions. Notably, while the data for the enzymes HincII (Fig. 6a) and XbaI (Fig. 6b) are well-described by a single strong protein-DNA interaction on either end of the protein, the data for the enzyme BsiWI (Fig. 6c) is not. Though the low-force protein signatures for BsiWI resemble those of the other two enzymes superficially, our theoretical model allows us to distinguish its unzipping behavior. In this case, the data are consistent with multiple weaker interactions surrounding a central, stronger region of contact.

**Figure 6:**
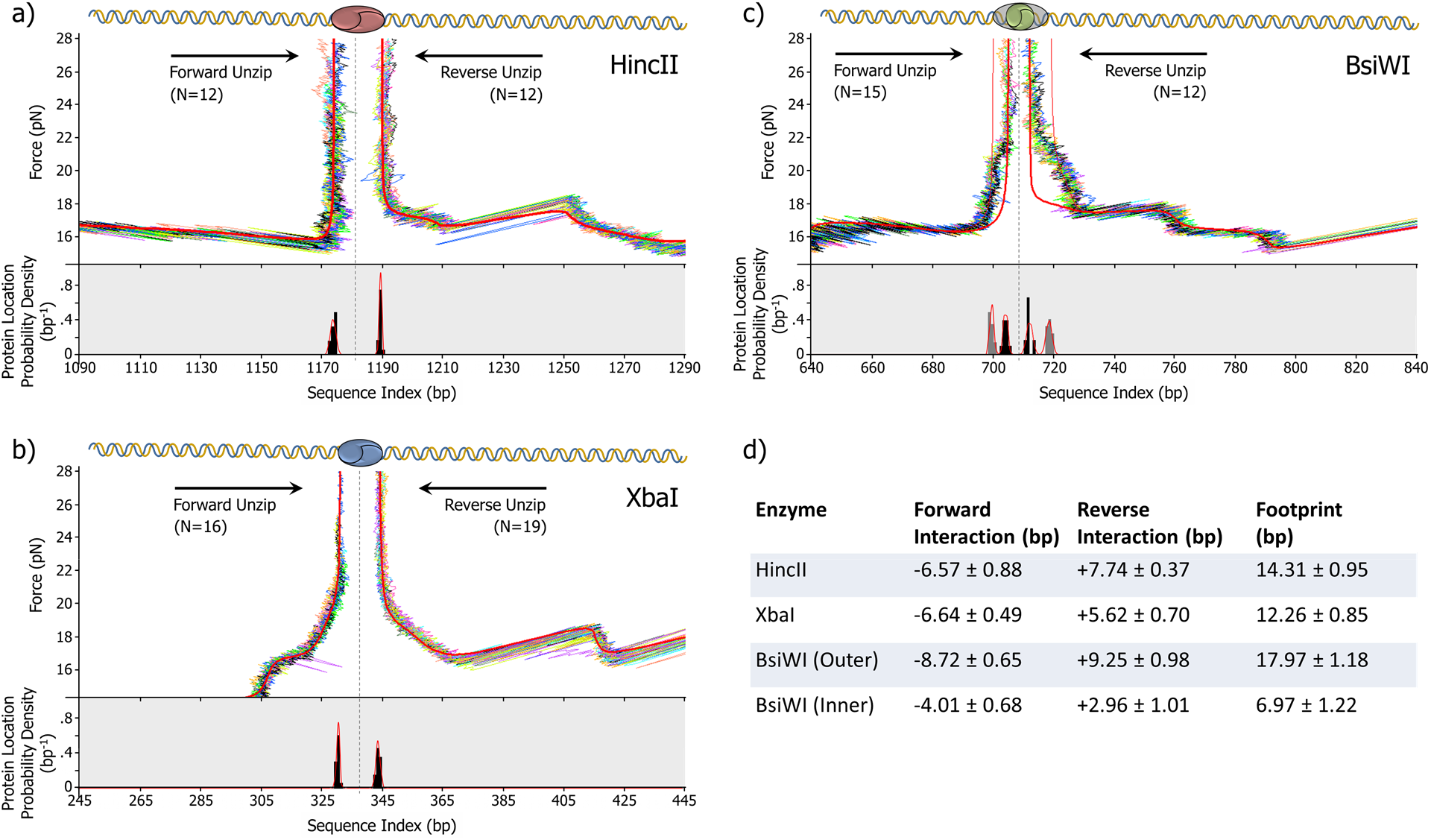
Restriction enzyme interaction mapping results. Three type II restriction enzymes, HincII **(a)**, XbaI **(b)**, and BsiWI **(c)**, were unzipped from either direction. For each trace (DNA molecule) we determined the most likely position of the protein using the *χ*^2^ technique from Figure 5c. A histogram of *χ*^2^ fitting results is shown below the data traces. The means and widths of the distributions were obtained using the maximum likelihood method. **(d)** A summary of the unzipping footprinting results. The precision of *χ*^2^ localization for individual interactions ranged from 0.37 bp to 1.01 bp.

As expected, the footprints of all three enzymes are centered with respect to their cut sites to within 1 bp, with sizes of 14.3 ± 1.0 bp (HincII), 12.3 ± 0.9 bp (XbaI), and 7.0 ± 1.2 bp (BsiWI). This is in good general agreement with previous DNAase I^30^ and unzipping^31^ footprint measurements on restriction enzymes of the same class, which have obtained footprints ranging from 5 to 20 bp for 8 different species. Of the three enzymes we unzipped, only HincII has a published crystal structure^32^. The protein shadows the bound DNA across a region of ^~^50 Å or 14.8 bp, within error of the 14.3 ± 1.0 bp we mapped by unzipping. The precision of the localization measurements is state-of-the-art; individual interactions were localized with standard deviations from 0.37 to 1.01 bp. This is the first reported instance of sub-bp precision in the localization of a protein by unzipping. These data show that, rather than being hindered by image tracking, dual optical trapping measurements can be enhanced by image tracking’s direct and robust measurement of extension in biological systems.

## Discussion

The photodiode-based technique of back focal plane interferometry (BFPI) has been widely acknowledged as the only detection method suitable for attaining the highest level of optical trapping performance for biophysical studies^33–35^. This has persisted in the field, even after advances in camera technology have improved the precision and speed of image tracking, due to the inability of cameras to measure the relative displacement of a bead from its trap center with the same accuracy and speed as BFPI, particularly in the important case of mobile traps.

By achieving an accurate and robust measure of force in an image-based optical trap, and by extending this ability to mobile optical traps without sacrificing accuracy or precision, we have developed an image-based optical trapping detection platform that yields state-of-the-art biological data. Thus equipped, we are now able to take full advantage of image tracking’s greatest strength: unlike BFPI, image tracking is in fact able to measure *extension* in a dual optical trap directly, making it potentially superior to other detection techniques in this aspect (Supplementary Fig. 5). This is evidenced by the exceptional quality of the instrument’s data, which has resulted directly in the elucidation of the mechanism by which a bound protein can affect the progression of an approaching unzipping fork.

With their compatibility with commercial microscopes and broad flexibility in implementation configurations, image-based techniques are familiar to biologists and biophysicists across a diverse array of applications. Along with these features, the added ability to realize the highest levels of performance in optical trapping detection renders image tracking an appealing option for making even the most demanding dual-trapping single-molecule measurements. This is particularly true in cases where BFPI is challenging to implement, e.g. with opaque substrates^36^, and with certain hybrid instruments^37^ where traditional optical trapping detection cannot be used. Ultimately, the fundamental limits of this detection method are set by the stability of the dual trap separation relative to the bandwidth and signal-to-noise ratio of the camera. Thus as the speed and noise characteristics of imaging technologies continue to improve at a rapid rate, image tracking may be poised to become the preferred detection method in dual optical traps.

## Methods

### Instrument Construction

A part list is located in Supplementary Note 1, and a full instrument layout is presented in Supplementary Figure 1. Dual optical traps are generated by timesharing a single 1064-nm laser at 50 kHz via an AOD. The traps are sourced from a 5W, 1064-nm fiber-coupled laser (IPG Photonics). A 60×, 1.27 NA, IR-corrected water immersion objective (Nikon) permits stable trapping deep within the sample chamber, facilitating DNA dumbbell tether formation and experimental flexibility. An acousto-optic deflector (AOD) (Gooch and Housego) is used to generate timeshared dual traps from the single CW laser source. Lenses map the beam rotation at the AOD onto the back focal plane of the microscope objective, and expand the beam to overfill the objective’s back aperture. The AOD is driven by an RF synthesizer (Gooch and Housego), from which frequencies between 35 and 45 MHz produce a trap displacement in the sample plane of up to about 10 μm. An FPGA (National Instruments 7852R) directs the output of the RF synthesizer. This “trap control FPGA” runs on a 40 MHz internal clock, enabling flexible timing for trap modulation and signal acquisition with a precision of 25 ns. We modulate the traps at 50 kHz, resulting in an “on” time for each trap of 10 μs.

The diffraction efficiency of an AOD is not constant with respect to the driving frequency. To produce traps of equal and uniform stiffness across their mobile range, we perform feedback on the trap powers (Supplementary Fig. 2). This requires a high-speed detector in order to measure each trap’s power individually during its portion of the modulation cycle. Silicon detectors are not well-suited to this purpose due to parasitic filtering at near-infrared wavelengths^38^, therefore we use a high-speed InGaAs photodiode (Thorlabs DET20C). Using the timing capabilities of the FPGA, a measurement sample of the analog photodiode output is acquired during the center of each trap’s on period, and then individual trap power feedback is performed on the FPGA to determine the trap output powers for the next modulation cycle.

To generate images for bead position detection, we use a high-speed CMOS camera (Mikrotron CAMMC1362) and illuminate the microscope sample from above with a high-powered LED (Thorlabs) emitting at 455 nm. The LED provides up to 5.4W of power, allowing high contrast and low-noise images even at 10,000 frames per second (fps) (Supplementary Fig. 3). A low-powered LED operating at 630 nm also illuminates the sample from above for wide-field-of-view imaging at 30 fps by an additional CCD camera (Andor), which in our case is a high-sensitivity camera which can also be used to acquire fluorescent images. This second camera primarily aids in tether formation and sample chamber navigation, and any basic CCD camera will suffice in this role. The two imaging wavelengths are combined for illumination using a dichroic mirror (Thorlabs). A second dichroic mirror (Chroma, 5 mm thick) below the objective transmits the trapping laser, and reflects visible wavelengths towards our imaging apparatus. The visible light is collected by a tube lens (Thorlabs), and then directed to their respective cameras by a third dichroic (Chroma). A 4× optical magnification attachment (Nikon) on the Mikrotron camera provides additional image enlargement. The conversion from camera pixels to nanometers in the sample plane is determined using a calibration grid printed on a microscope slide (Thorlabs), from which we obtained a value of 57.3 nm per pixel (0.01745 ± 0.00001 pixels per nm) using a cross-correlation-based analysis of the grid images.

### FPGA Image Tracking

Image tracking is performed on a second “image tracking FPGA” (National Instruments 1473R). The high-speed camera is connected directly to the FPGA via CameraLink in the base configuration. Pixels from a cropped region of interest 200×7 pixels in size are sent in single tap, 10-bit mode on the camera’s 80 MHz clock. On the image tracking FPGA, three parallel, independent loops perform image acquisition and pre-processing, bead position tracking, and transfer of the tracked bead positions, respectively (Fig. 2a). In the acquisition loop, pixels are read out one at a time and processed as they arrive. Sequential processing steps are pipelined to allow for full parallelism; once a pixel leaves an individual processing step to the next downstream function, a new pixel is clocked in, so that all functions are acting on a different pixel simultaneously.

Pixel pre-processing begins by applying a correction for the camera sensor’s fixed pattern pixel noise by subtracting a constant offset for each pixel. The appropriate offset for each pixel is determined before an experiment by averaging 1,000 image frames and calculating each pixel’s difference from the mean frame intensity, and then sent and stored in the FPGA’s block memory. Next, the pixel stream is routed to one of two memory queues, one for each bead’s pre-defined sub-ROI within the full image ROI. As the pixels for a given frame are being sorted into memory, the FPGA determines the *x* and *y* indices of the pixel with the maximum brightness value for each bead’s sub-ROI, and checks the max pixels’ location and brightness value against user-defined validation conditions to determine if each sub-ROI contains a valid, trackable bead. This maximum brightness pixel in each bead’s sub-ROI provides a coarse estimate of the bead center about which the tracking algorithm can later focus. On the last pixel of an image frame, the validation results for each bead’s max pixel is passed to the FPGA tracking loop running on a 40 MHz clock.

The tracking loop is triggered to begin once it receives the validation results from the acquisition loop. Pixels are read out of their respective queues for each bead sequentially. If the frame contains a valid bead, the tracking loop first constructs a line profile centered around the maximum brightness pixel identified in the acquisition loop. The size of the line profile may be varied, but for this work we used a line profile 15 pixels in length, summing across three pixels in the orthogonal direction to reduce noise. The line profile is zeroed by subtracting the value of the first element of the array from all array elements, which greatly reduces tracking bias. We then cross-correlate the zeroed line profile with its mirror image to generate a one-dimensional cross-correlation array, the peak of which can be mapped back to the original pixel space, giving the computed location of the bead. The sub-pixel maximum of the cross-correlation array is found by performing a parabolic least squares fit to the array maximum and its single nearest neighbor to either side. FPGAs are not generally well-suited to nonlinear curve fitting, however, the task is greatly simplified in this case by the fact that the Vandermonde Matrix and its transpose are fixed and can be calculated in advance. For a least squares fit about the cross-correlation array peak value *R*_*i*_ and its nearest neighbors *R*_*i*−1_ and *R*_*i*+1_, the maximum of the resulting curve fit reduces to the following “three point estimator”:

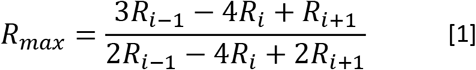

Because multiply and divide operations performed on an FPGA require specialized hardware multipliers of which an FPGA typically has just a few dozen, the ability to simplify a curve fitting process involving matrix calculations and determinants to a single, simple arithmetic expression is crucially important for the viability of performing this image processing algorithm on an FPGA

The pixel locations of each bead determined in the tracking loop are passed to the data transfer loop, which sends these values to the trap control FPGA. The two FPGAs are connected via a dedicated I/O extension board (National Instruments), which equips the image tracking FPGA with 8 configurable digital input/output (DIO) lines which we wired to 8 of the DIO lines on the trap control FPGA. The image tracking FPGA connects to the I/O extension board directly via a ribbon cable; signals are not routed through the computer’s PCIe bus. The tracked pixel locations of the two beads are stored in a 30-bit fixed point data type yielding a discretization of 2.38×10^-7^ pixels or 1.37×10^-5^ nm, which are here encoded in a length-30 Boolean array suitable for digital TTL transfer. We use one DIO line to signal the trap FPGA that data are ready for transfer, a second to clock the data transfer at 10 MHz, and the remaining 6 DIO lines to transfer the 30-bit bead positions over 5 clock cycles. The full transfer of one bead position data point takes 800 nanoseconds. The data are read by the trap control FPGA and translated back into a numeric form, at which point the current AOD frequency of the active trap is used to index the trap separation LUT for calculatingΔ*X*_bead_. The latency of bead tracking is thus a critical parameter for this technique, as the bead separation data lags the trap separation data by the tracking latency time which will result in force errors when the traps are in motion. Because our tracking is implemented on an FPGA, the total latency between the finished camera exposure and the completed transfer of the tracked bead positions is around 50 μs, yielding errors less than an Angstrom in Δ*X*_bead_ for active trap speeds up to 400 nm/s.

### Free Bead Separation Calibration and Δ*X*_bead_ Determination

To calibrate the free bead separation, we start with an untethered bead in each trap. The stationary trap remains fixed at 35 MHz and the active trap is stepped from 37 to 45 MHz across 330 intervals spaced evenly in frequency as the bead separation is measured. The spacing of the trap separation LUT, resulting in one data point approximately every 22 nm of trap movement, was determined empirically to maximize the accuracy of Δ*X*_bead_. At each LUT position, data were collected for 200 ms and then averaged. The array of trap separations is transferred to the trap control FPGA and stored in block memory. During an experiment, upon arrival of tracked bead positions from the image tracking FPGA, the bead-to-bead separation is calculated and the current position of the active trap is used to index the trap separation LUT with linear interpolation.

### Stretching and Unzipping Experiments

The generation of stretching and unzipping templates has been described previously^25,39,40^. For forming DNA dumbbell tethers, carboxylated beads (Polysciences) were coated with (either) streptavidin and anti-digoxigenin in house. The process of forming a DNA dumbbell tether is described in detail in Supplementary Figure 7.

For Figure 4, our template for both stretching and unzipping consisted of 6644 bp of double-stranded DNA “arms” and an 832-bp unzipping “trunk”. Experiments were performed in phosphate-buffered saline (10 mM Na_2_HPO_4_, 1.8 mM KH_2_PO_4_, pH 7.4, 137 mM NaCl, 2.7 mM KCl). In Figure 4a, the beads were 500 nm in diameter and the trap stiffness was constant at ^~^0.28 pN/nm. The active trap was moved away from the stationary trap to stretch the DNA arms at a constant velocity of ^~^200 nm/s. Data traces were truncated at a force of 14 pN, just prior to the onset of unzipping. In Figure 4b, the beads were 800-nm in diameter and the trap stiffness was constant at ^~^0.36 pN/nm. Unzipping was performed using a constant active trap velocity of 20 nm/s, and rezipping was performed using a constant active trap velocity of -40 nm/s.

Protein unzipping experiments for data shown in Figures 5 and 6, as well as all related calibration experiments, were performed in a buffer containing 50 mM Tris-HCL pH 7.9, 100 mM NaCl, and 1 mM CaCl_2_. Unzipping templates consisted of 6644 bp of double-stranded DNA arms and unzipping trunks of various lengths between 0.7 and 4.4 kbp. All experiments were performed under conditions of constant trap stiffness (ranging from 0.27 to 0.38 pN/nm) and constant active trap velocity (ranging from 40 to 100 nm/s). Proteins were purchased from New England Biolabs. One microliter of stock protein corresponding to 10 (HincII, BsiWI) or 20 (XbaI) vendor-calibrated units was added to a 15 μL pre-incubation of DNA and beads just prior to introduction into the experimental flow cell for tether formation. Additionally, protein was included in the unzipping buffer in which unzipping experiments were performed, at concentrations ranging from ^~^100 Units/mL (1:100 dilution) to ^~^700 Units/mL (1:15 dilution)

### Data Analysis

While force in an optical trap is generally treated as being linearly proportional to bead displacement, the validity of this approximation decreases with increasing distance from the trap center. To improve data accuracy throughout the wide range of forces encountered during protein unzipping experiments, we developed a technique to measure the non-uniformity of trap stiffness as a function of bead displacement. This calibration, performed once, is used to convert all measured bead displacements into forces in later experiments. The technique utilizes the knowledge that, in our constant trap velocity unzipping experiments, a DNA molecule will unzip at the same force regardless of the nature of the trapping potential.

Accordingly, we unzipped DNA molecules multiple times at laser trapping powers ranging from 120 to 300 mW in the sample plane, resulting in average bead displacements during unzipping of 117 to 33 nm. From our theoretical model of unzipping we obtained the predicted unzipping force, and plotted this force per watt of trapping power versus the measured bead displacement at each power (Supplementary Fig. 8). The resulting data was well-fit by the derivative of a Gaussian (DoG) function. This function and the obtained fit parameters were subsequently used to convert measured bead displacements into values of force. For protein unzipping data only, we allowed the amplitude of the DoG function to be a free parameter in order to correct for small observed variations in unzipping force (typically 1 to 3%) between traces, which could be due, e.g., to variations in bead sizes or drift of the trapping power in the sample plane. The unzipping force calculated by our theoretical model was used to determine the appropriate DoG amplitude for the conversion of each trace.

Thusly determined values of force and extension during unzipping were converted to the number of bases unzipped in the sequence using a modified Marko-Siggia model^21^ for double-stranded DNA (dsDNA), and the freely-jointed chain model^41^ for single-stranded DNA (ssDNA). DNA elasticity parameters may be obtained by fitting double and single-stranded DNA stretching data with their respective models. For this work, we performed this calibration for ssDNA and used previous calibration results^21^ for dsDNA (persistence length of 42 nm, elastic modulus of 1200 pN, and contour length per base of 0.338 nm, also used for stretching theory curve in Figure 4a). To stretch single-stranded DNA, a hairpin was placed at the end of an unzipping segment, and the construct was unzipped to completion and then stretched. From the resulting values of extension, the contribution of the double-stranded portion of the unzipping construct was subtracted, leaving the force-extension relationship of only the single-stranded portion for fitting. Fitting resulted in an ssDNA persistence length of 0.71 nm (half the Kuhn length), elastic modulus of 420 pN, and counter length per base of 0.52 nm. Once represented as force vs the number of base pairs unzipped, individual protein unzipping traces were aligned horizontally to the DNA baseline theory using small global shift and stretch adjustments (typically < 10 nm and < 1%, respectively) determined by a simplex algorithm.

Around the approximate apparent protein position as indicated by the location of the vertical force rise asymptote, we generated theory curves for proteins located at integer-spaced bp locations along the template. The theory assumes an infinite energy barrier at the specified point along the sequence, such that the unzipping fork cannot progress further. For each generated theory curve, we computed the χ2 value comparing our measured base pairs unzipped to the theoretical base pairs unzipped, using DNA extension as the common independent variable. To determine protein location with sub-bp precision, we plot the resulting *χ*^2^ vs theoretical protein position and fit the minimum point plus its nearest neighbor to either side with a parabola.

The footprints for HincII and XbaI were determined using *χ*^2^ fitting on the full unzipping signature, from the onset of the deviation from the naked DNA theory to a force of 26 pN. To analyze the BsiWI footprint, we performed *χ*^2^ fitting on cropped regions of the protein unzipping signature, from the onset of the force deviation from the naked DNA to a force of 20 pN for the outer interaction, and from 23 to 26 pN for the inner interaction.

All analysis steps were performed with custom LabVIEW 2015 software.

## Supplementary Information

Additional data, instrument details, and technical information are contained in the supplementary materials available for download.

## Acknowledgements

We thank the members of the Wang Laboratory for critical discussion and comments on this work. We especially thank Drs. Tung Le and Fan Ye for aiding in DNA template construction, Dr James Baker for coating beads, and Ryan Badman for SEM imaging of beads. We thank Professor Matthew Comstock for discussion and example software related to the timesharing component of our instrument. This work was supported by the Howard Hughes Medical Institute, the National Institutes of Health (T32GM008267), and the National Science Foundation (MCB-0820293 to M.D.W. and a Graduate Research Fellowship DGE1144153 to J.L.K.).

